# Hypoxia induces transcriptional and translational downregulation of the type I interferon (IFN) pathway in multiple cancer cell types

**DOI:** 10.1101/715151

**Authors:** Ana Miar, Esther Arnaiz, Esther Bridges, Shaunna Beedie, Adam P Cribbs, Damien J. Downes, Robert Beagrie, Jan Rehwinkel, Adrian L. Harris

## Abstract

Hypoxia is a common phenomenon in solid tumours and is considered a hallmark of cancer. Increasing evidence shows that hypoxia promotes local immune suppression. Type I IFN is involved in supporting cytotoxic T lymphocytes by stimulating the maturation of dendritic cells (DCs) and enhancing their capacity to process and present antigens. However, there is little information about the relationship between hypoxia and the type I interferon (IFN) pathway, which comprises the sensing of double-stranded RNA and DNA (dsRNA/dsDNA), followed by IFNα/β secretion and transcription activation of IFN-stimulated genes (ISGs). The aims of this study were to determine both the effect and mechanisms of hypoxia on the I IFN pathway in breast cancer.

There was a downregulation of the type I IFN pathway expression at mRNA and protein level in cancer cell lines under hypoxia *in vitro* and *in vivo* in xenografts. This pathway was suppressed at each level of signalling, from the dsRNA sensors (RIG-I, MDA5), the adaptor (MAVS), transcription factors (IRF3, IRF7, STAT1) and several ISGs (RIG-I, IRF7, STAT1, ADAR-p150). There was also lower IFN secretion under hypoxic conditions. HIF1 and HIF2 regulation of gene expression did not explain most of the effects. However, ATAC-Seq data revealed that in hypoxia peaks with STAT1 and IRF3 motifs had decreased accessibility.

Thus hypoxia leads to an overall 50% downregulation of the type I IFN pathway due to repressed transcription and lower chromatin accessibility in a HIF1/2α-independent manner, which could contribute to immunosuppression in hypoxic tumours.

## INTRODUCTION

Oxygen can only diffuse 100-180μm from the nearest capillary and consequenctly, poorly vascularized tumours or those that grow quickly suffer from hypoxia, low pH and nutrient starvation. Hypoxia is closely linked to the hallmarks of cancer as it induces the switch to glycolytic metabolism, promotes proliferation, enhances resistance to apoptosis, induces unlimited replication potential and genomic instability, reduces immuno-surveillance, and induces angiogenesis and migration to less hypoxic areas (1).

This hypoxia response is driven by a family of dimeric hypoxia-inducible transcription factors (HIF-1, HIF-2, HIF-3) which are composed of an oxygen-sensitive α-subunit (HIF-1α, HIF-2α, HIF-3α) and a constitutively expressed β-subunit (HIF-1β) (2). Under well oxygenated conditions, HIF-1/2α are bound by the von Hippel-Lindau (VHL) protein which targets them for proteasomal degradation (3). VHL binding is dependent upon hydroxylation of specific proline residues by prolyl hydroxylase PHD2, which uses O2 as a substrate, and as a consequence its activity is inhibited under hypoxia (4).

Hypoxia induces an immunosuppressive microenvironment; it inhibits the ability of macrophages to phagocytose dead cells, present antigens to T cells, inhibits the anti-tumour effects of macrophages (5), and favours their differentiation to M2 type which is more immunosuppressive (6). T-cell-mediated immune functions are downregulated by low oxygen levels, resulting in impairment of cytokine expression and less T-cell activation (7). Moreover, the cytotoxic potential of natural killer (NK) cells is reduced under hypoxia resulting in a decreased anti-tumour response that allows metastasis (8). Type I interferons (IFNs) comprise multiple IFNα’s (encoded by 13 genes), IFNβ (encoded by one gene) and other less studied IFNs (IFNε, IFNκ, IFNω). They are produced by most cell types and their transcription is upregulated after the activation of pattern recognition receptors (PRR) which sense viral/bacterial dsRNA (through RIG-I, MDA5, or TLRs) and dsDNA/unmethylated DNA (through cGAS, DDX41, IFI16, or TLRs). Type I IFNs bind to the heterodimeric IFNα/β receptor composed of IFNAR1 and IFNAR2, triggering the JAK-STAT pathway and activating the transcription of IFN-stimulated genes (ISGs) (9) (Supplementary figure 1). Type I IFNs are involved in all three phases of the cancer immunoediting process by which the immune system eliminates the malignant cells, moulds the immunogenic phenotypes of developing tumours and selects those clones with reduced immunogenicity (10). Type I IFNs support cytotoxic T lymphocytes by stimulating the maturation of dendritic cells (DCs) and enhancing their capacity to process and present antigens (11). Endogenous IFNα/β is required for the lymphocyte-dependent rejection of highly immunogenic sarcomas in immunocompetent hosts (10), and they are crucial for the innate immune response to transformed cells as *Ifnar1^−/−^* CD8^+^ dendritic cells are deficient in antigen cross-presentation, and fail to reject highly immunogenic tumours (12). Moreover, type I IFNs boost immune effector functions by enhancing perforin 1 and granzyme B expression (13), and promoting survival of memory T cells.

Type I IFN is involved in the success of current anticancer treatments both directly (tumour cell inhibition) and indirectly (anti-tumour immune responses to the nucleic acids and proteins released by dying cells, recently named immunogenic cell death, ICD) (14) (9). In mouse models, IFNAR1 neutralization using antibodies blocked the therapeutic effect of antibodies against human epidermal growth factor receptor 2 (HER2) or epidermal growth factor receptor (EGFR) (15, 16). Moreover, radiotherapy, via DNA damage, induces intratumoural production of IFNβ which enhances the cross-priming capacity of tumour-infiltrating dendritic cells (17). Chemotherapeutic agents such as anthracyclines, bleomycin or oxalaplatin induce ICD, and the activation of an IFN-dependent program is required for successful therapy (18, 19).

Tumour hypoxia creates resistance to many cancer treatments as oxygen is essential for ROS formation to kill cells during ionizing radiation (20, 21), and HIF1α upregulation mediates resistance to chemotherapy by upregulating anti-apoptotic and pro-survival genes (22).

Thus we investigated the regulation of the type I IFN pathway under hypoxia at basal levels and upon activation of IFN signalling using the dsRNA mimic polyinosinic:polycytidylic acid (poly I:C). We found there was substantial HIF-independent downregulation of IFN signalling which could affect all the before mentioned therapies and contribute to treatment resistance.

## MATERIAL AND METHODS

### Cell culture and transfection

All cell lines used are listed in supplementary table 1. They were cultured in DMEM low glucose medium (1g/l) supplemented with 10% FBS no longer than 20 passages. They were mycoplasma tested every 3 months and authenticated during the course of this project. Cells were subjected to 1% or 0.1% hypoxia for the periods specified in each experiment using an InVivO2 chamber (Baker).

Transfection of poly I:C (P1530, Sigma) was performed in Optimem reduced serum (11058021, Thermo Fisher Scientific) medium for 6h unless it is otherwise indicated. Lipofectamine 2000 (11668019, Thermo Fisher Scientific) was used following the manufacturer’s instructions.

### Western blot and RT-qPCR

Methods detailed in Supplementary materials.

### IFN bioassay

Performed as previously described (23) and detailed in Supplementary Materials.

### Single Cell sequencing and analysis

Single-cell RNA-seq libraries were prepared as per the Smart-seq2 protocol by Picelli et al (24) with minor technical adaptations detailed in Supplementary material. scRNA-seq data were deposited in Gene Expression Omnibus under SuperSeries accession number GSE134038 and detailed in Supplementary materials.

### Xenograft growth and IHC

Procedures were carried out under a Home Office license. Xenograft experiments were performed as described in (25) and detailed in Supplementary materials.

### ATAC-seq and analysis

Samples were prepared and analysed as previously described (26) with minor modifications detailed in Supplementary materials. ATAC-seq data is available in the Gene Expression Omnibus (GSE133327).

## RESULTS

### Hypoxia causes downregulation of the type I IFN pathway in unstimulated cells both at mRNA and protein level

MCF7 cells were subjected to normoxia, 1% hypoxia or 0.1% hypoxia for 48h. This lead to a decrease of both protein (figure 1A) and RNA (figure 1B) of members of the type I IFN pathway. The effect was seen from dsRNA sensors (RIG-I [encoded by *DDX58*], MDA5 [encoded by *IFIH1*]), adaptor MAVS, transcription factors triggering IFNα/β production (IRF3, IRF7) and transcription factors involved in ISGs’ activation (STAT1, STAT2), to downregulation of downstream ISGs (exemplified by ADAR-p150). This inhibitory effect was oxygen concentration-dependent and 0.1% hypoxia caused a greater downregulation of the pathway than 1% hypoxia.

**Figure 1.**
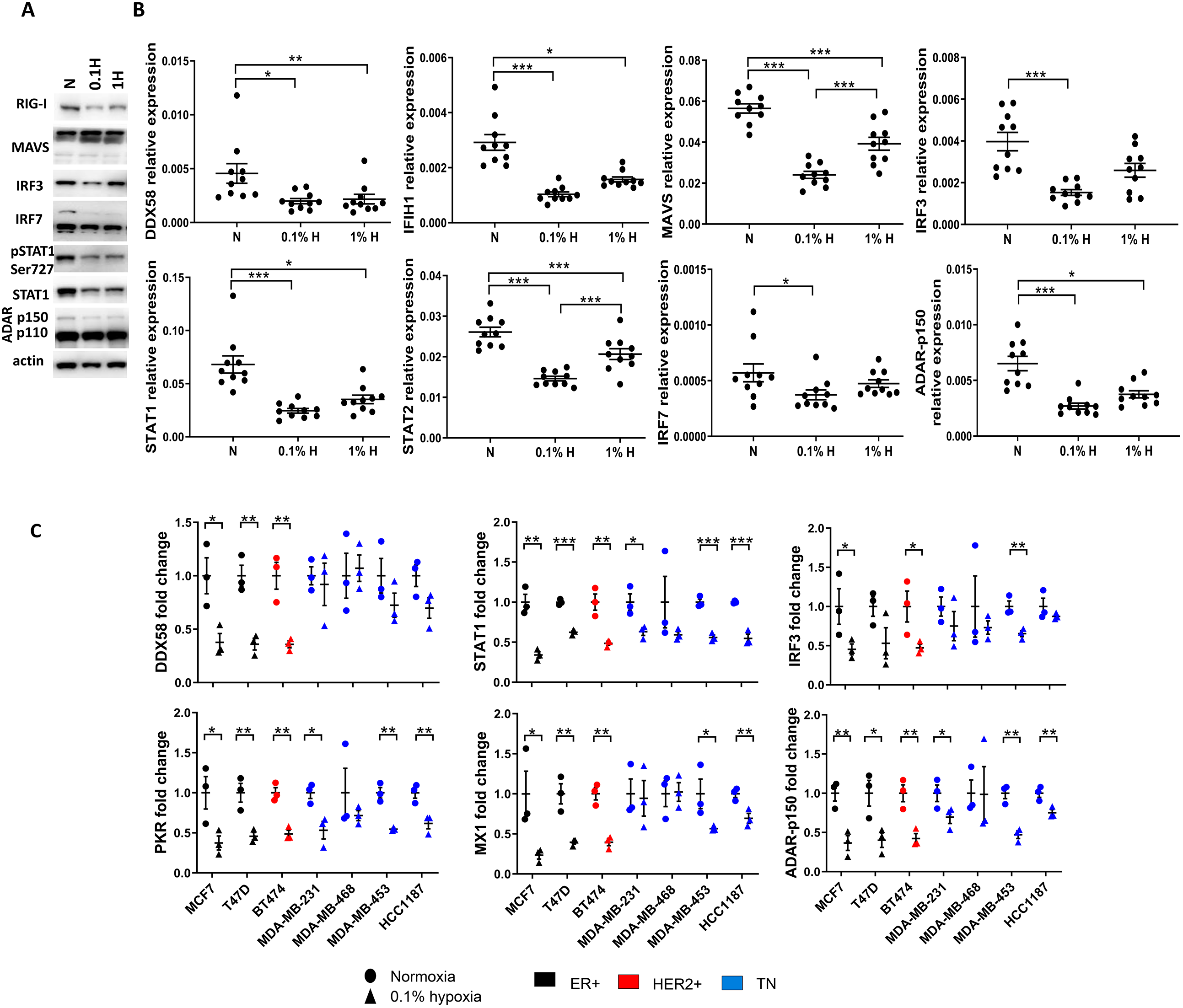
Type I interferon (IFN) pathway expression in breast cancer cells. A) Western-blot showing protein levels of different genes involved in the pathway in MCF7 cells cultured in normoxia, 0.1% and 1% hypoxia for 48h. B) qPCR data showing mRNA expression of genes involved in the pathway in the same experiment (n=10). C) mRNA expression of genes in the type IFN pathway in different breast cancer cell lines cultured in normoxia or 0.1% hypoxia for 24h (n=3). * p<0.05, ** p<0.01, *** p<0.001.

This finding was confirmed in other breast cancer types including oestrogen receptor positive (ER+; T47D), HER2+ (BT474) and triple negative (TN; MDA-MB-453, MDA-MB-231, HCC1187, MDA-MB-468) cells. In general, a downregulation of the type I pathway was observed in all cell lines in hypoxia. However, TN cell lines seemed to be more resistant to the effect of hypoxia, showing a lower or no downregulation of some transcripts such as *DDX58*, *IRF3* or *MX1* (figure 1C).

### Single cell transcriptomic analysis of the effects of hypoxia on the type I IFN pathway

Next, we performed a single cell RNA-Seq experiment using MCF7 cells subjected to 0.1% hypoxia for 72h. We found that most of the genes in the type I IFN pathway (such as *DDX58, IRF3* or *MX1*) are significantly less expressed in hypoxia (figure 2A).

**Figure 2.**
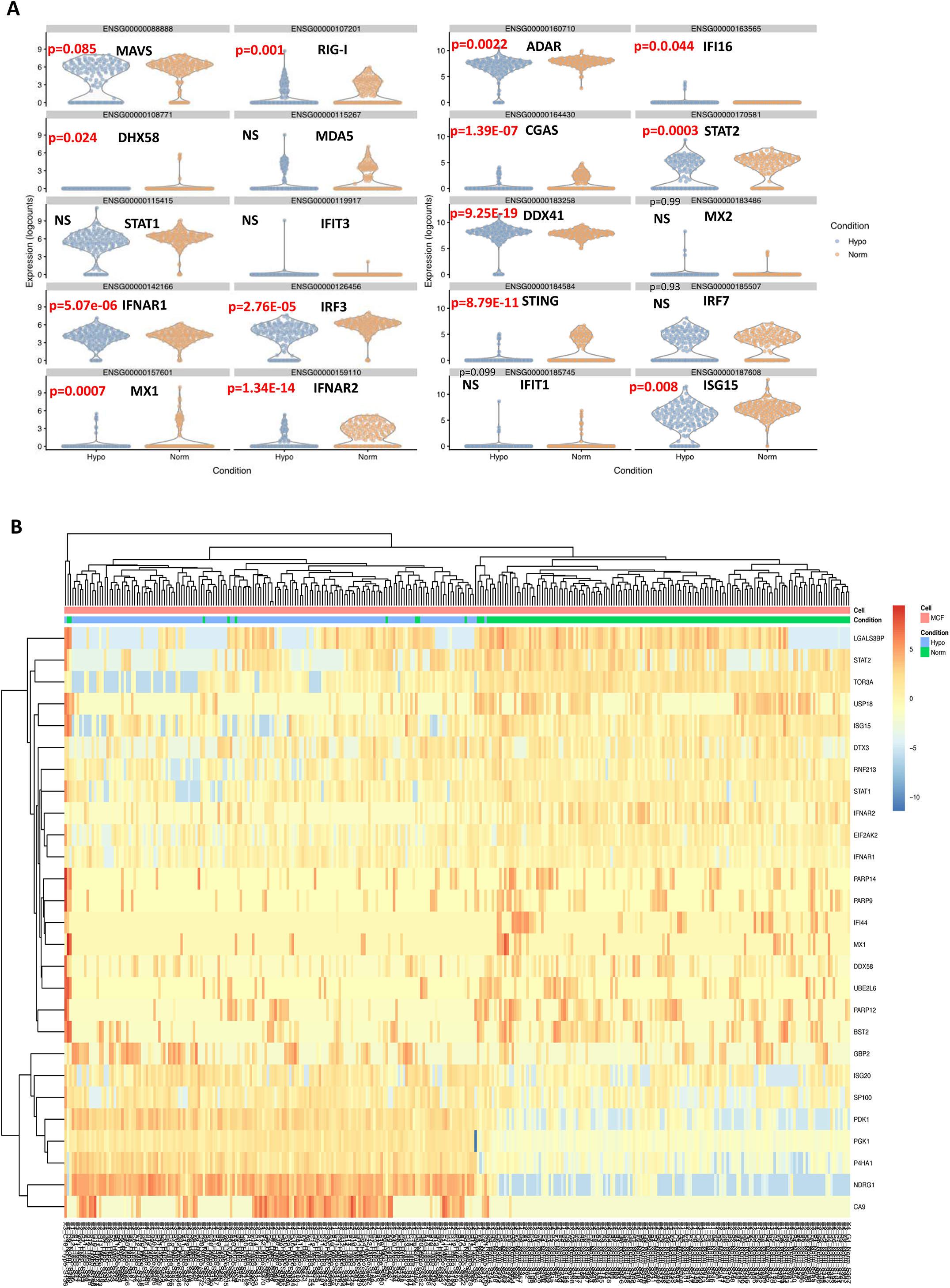
Single-cell analysis of type I IFN pathway in MCF7. A) Violin plots showing expression (log counts) of type IFN genes in normoxia (orange) and 0.1% hypoxia (blue) after 72h. B) Heatmap correlating mRNA expression type I IFN genes previously published with hypoxia target genes.

We also observed a decrease in the expression of several genes associated with the DNA sensing branch (*IFI16, TMEM173* [encodes STING], *DDX41* and *MB21D1* [encodes cGAS]) in MCF7 cells exposed to hypoxia. Next, we investigated the expression of a number of well described ISGs(27). This revealed a clear subpopulation of cells in which ISGs were downregulated following exposure to hypoxic conditions, e.g. *DDX58, IFIH1, STAT1, STAT2, IFIT1, IFIT3* or *MX2*. In contrast, HIF1α targets such those involved in glycolysis, were upregulated as expected (*PDK1, PGK1, P4HA1, NDRG1, CA9;* figure 2B).

### Reoxygenation after hypoxia leads to recovery of the type I IFN pathway

A time course in 0.1% hypoxia up to 48h showed that type I IFN downregulation at protein level was obvious at 48h in hypoxia (figure 3A), whereas at RNA level, there was a significant decrease from 16h onwards in *IFIH1, MAVS, STAT1* and *ADAR* (figure 3B). MCF7 cells were incubated in normoxia or 0.1% hypoxia for 48h and subjected to reoxygenation for 8h, 16h or 24h. At protein level (figure 3C), hypoxia caused the downregulation of the pathway but reoxygenation up to 24h did not reverse this effect except for RIG-I. However, at RNA level (figure 3D), reoxygenation caused a gradual upregulation of type I IFN pathway levels. Thus RNA expression responded more rapidly to O_2_ tension than protein.

**Figure 3.**
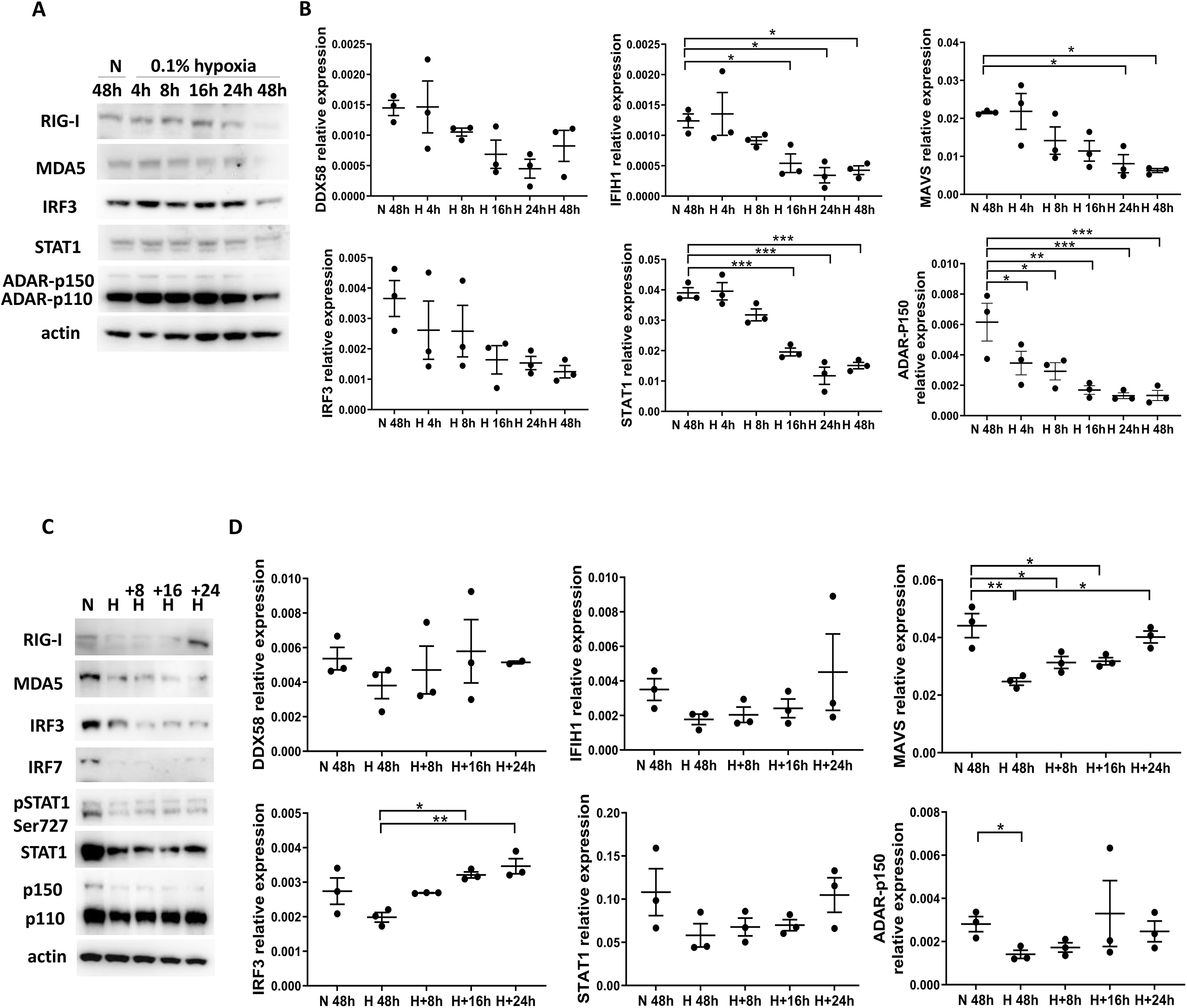
Type I IFN over time and during reoxygenation in MCF7 cells. A) Western-blot showing protein levels of different genes involved in the pathway in MCF7 cells cultured in normoxia for 48h or 0.1% hypoxia for 4h, 8h, 16h, 24h and 48h. B) qPCR data showing mRNA expression of genes involved in the pathway in the same experiment (n=3). C) Western-blot showing protein changes in MCF7 when exposed to normoxia or 0.1% hypoxia for 48h and after reoxygenation for 8h, 16h and 24h. D) qPCR data showing mRNA expression changes for the same experiment (n=3). * p<0.05, ** p<0.01, *** p<0.001.

### Hypoxia downregulates the type I IFN pathway at protein level upon stimulus with a dsRNA mimetic

MCF7 cells were incubated in normoxia or 0.1% hypoxia for 48h and transfected with poly I:C (mimetic of dsRNA) in the last 6h of the incubation to determine if the downregulation caused by hypoxia would persist upon activation. Poly I:C activated the type I IFN pathway as shown using pIRF3-Ser386 and pSTAT1-Tyr701; however hypoxia led to lower activation (lower levels of pIRF3 and pSTAT1) and to lower expression of RIG-I, MDA5, MAVS and IRF7 (figure 4A). These findings suggested that production/secretion of IFNs was lower in hypoxia than in normoxia.

**Figure 4.**
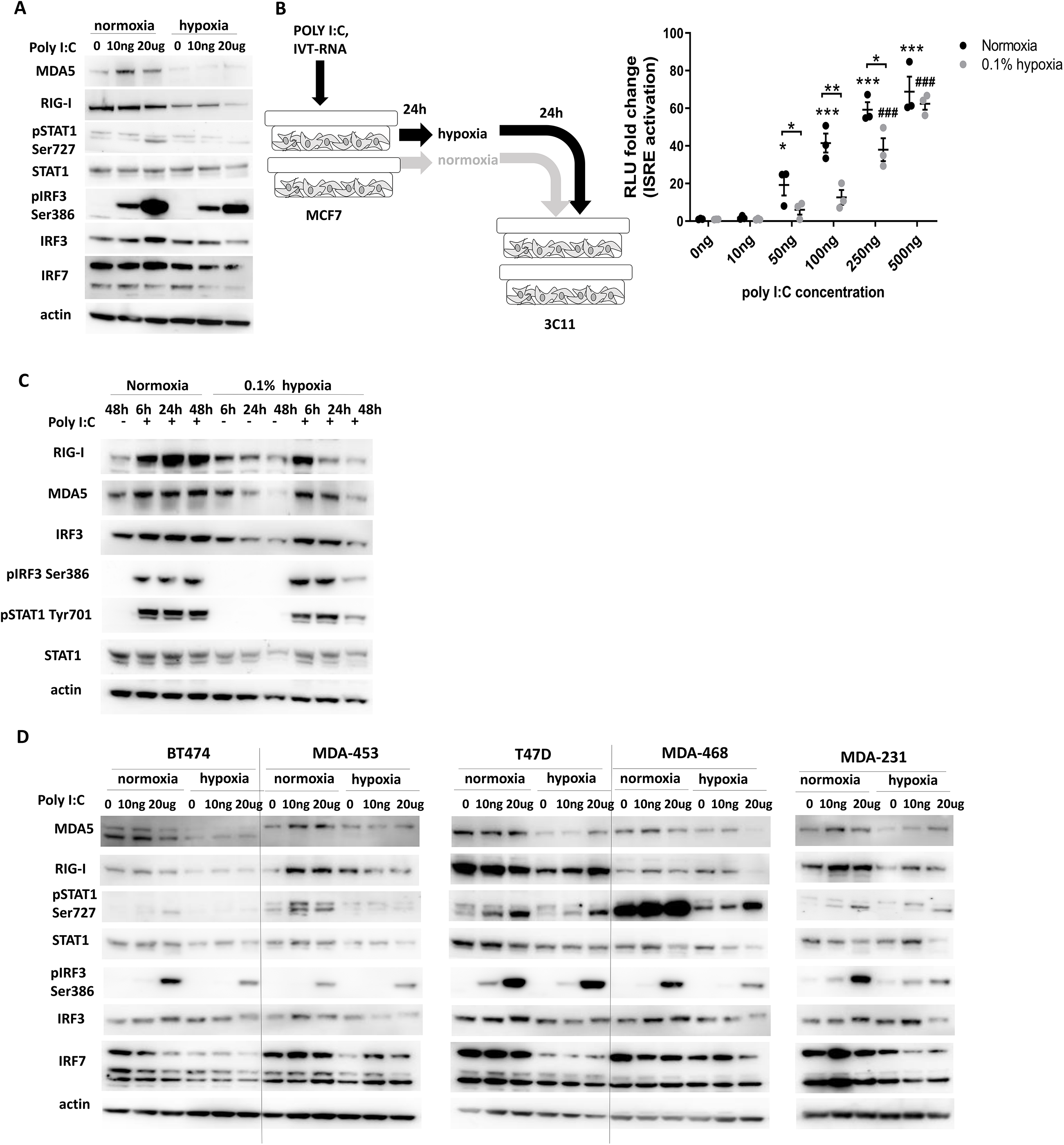
Type I IFN activation by poly I:C in breast cancer cells. A) Western-blot showing protein levels of MCF7 cells cultured in normoxia or 0.1% hypoxia for 48h and transfected with poly I:C in the last 6h. B) IFN bioassay showing Interferon Stimulated Response Element (ISRE) activation via luciferase activity in HEK293 cells using supernatants of MCF7 cells previously treated with normoxia or 0.1% hypoxia for 24h and transfected with different concentrations of poly I:C (n=3). C) Western-blot showing protein changes in different breast cancer cells when exposed to normoxia or 0.1% hypoxia for 48h and transfected with 20ug/ml poly I:C in the last 6h of the treatment. * p<0.05, ** p<0.01, *** p<0.001 in normoxia and normoxia vs hypoxia, # p<0.05, ## p<0.01, ### p<0.001 in hypoxia samples.

To investigate this, the stimulation of IFN-stimulated response element (ISRE) by potential IFNs present in the supernatant from normoxic or hypoxic cells was measured using indicator cells (HEK293-3C11, hereafter 3C11) (23). 3C11 cells harbour a stably integrated luciferase reporter under control of an ISRE, allowing the measurement of type I IFNs in supernatant samples. 3C11 cells were incubated with supernatant from MCF7 cells previously treated with different concentrations of poly I:C and normoxia or hypoxia, as described in the Supplementary Materials section. At all poly I:C concentrations, hypoxic supernatants showed lower activation of ISRE confirming the lower production/secretion of IFNs in these conditions (figure 4B).

A time course in hypoxia was performed using MCF7 cells cultured for 6h, 24h and 48h in normoxia or 0.1% hypoxia and transfected with poly I:C in the last 6h of incubation. pIRF3 and pSTAT1 were induced upon poly I:C transfection and their expression was downregulated when the cells were exposed to 48h hypoxia but not at shorter incubations. RIG-I, MDA5 and MAVS were downregulated at 24h, suggesting that hypoxia diminishes the sensitivity of this pathway, particularly effecting late signalling in the amplification phase (figure 4C).

The reponse to hypoxia was analysed in a panel of breast cancer cell lines which were incubated in normoxia or 0.1% hypoxia for 48h and transfected with poly I:C during the last 6h. Again, the activation of the pathway was lower in hypoxia and the expression of MDA5, RIG-I, STAT1 and IRF7 was also decreased, showing a general effect in breast cancer cell lines (figure 4D).

We assessed cell lines from other cancer types or normal tissues e.g. HepG2 hepatocarcinoma, HKC8, a non-transformed proximal renal tubular cell line and fibroblasts. The pSTAT1-Tyr701 response to poly I:C was reduced in all cell lines in hypoxia (supplementary figure 2). Interestingly, pIRF3 expression in HepG2 was higher in hypoxia than in normoxia compared to IRF3 expression which was lower. This different response may contribute to the higher viral replication under hypoxia in hepatocarcinoma (28). In HKC8 cells and fibroblast, IRF3 expression was higher in hypoxia or did not change but pIRF3 was reduced. Thus, hypoxia causes a general downregulation of the type I IFN pathway in cancer, and similarly in normal cell lines.

### Type I IFN downregulation is partially dependent on HIF1α

To determine the role of HIF1α in the results observed under hypoxic conditions, we used MCF7-HIF1α-KO and MCF7-WT cells. In MCF7-WT, hypoxia suppressed RIG-I, MDA5 and IRF3 expression. Interestingly, in MCF7-HIF1α KO cells in normoxia, RIG-I and STAT1 levels were higher than in MCF7-WT cells, supporting a partial role for HIF1α in suppressing them. However, in hypoxia, STAT1 and RIG-I levels were reduced, indicating a HIF1α-independent mechanism to further regulate them maybe compatible with residual HIF2 effect.

Poly I:C in both cell lines in normoxia induced IRF3, pIRF3, IRF7 and pSTAT1. This induction was higher in MCF7-HIF1α-KO cells, apart from pSTAT1, which was still induced but to lower level than the MCF7-WT cells (figure 5A).

**Figure 5.**
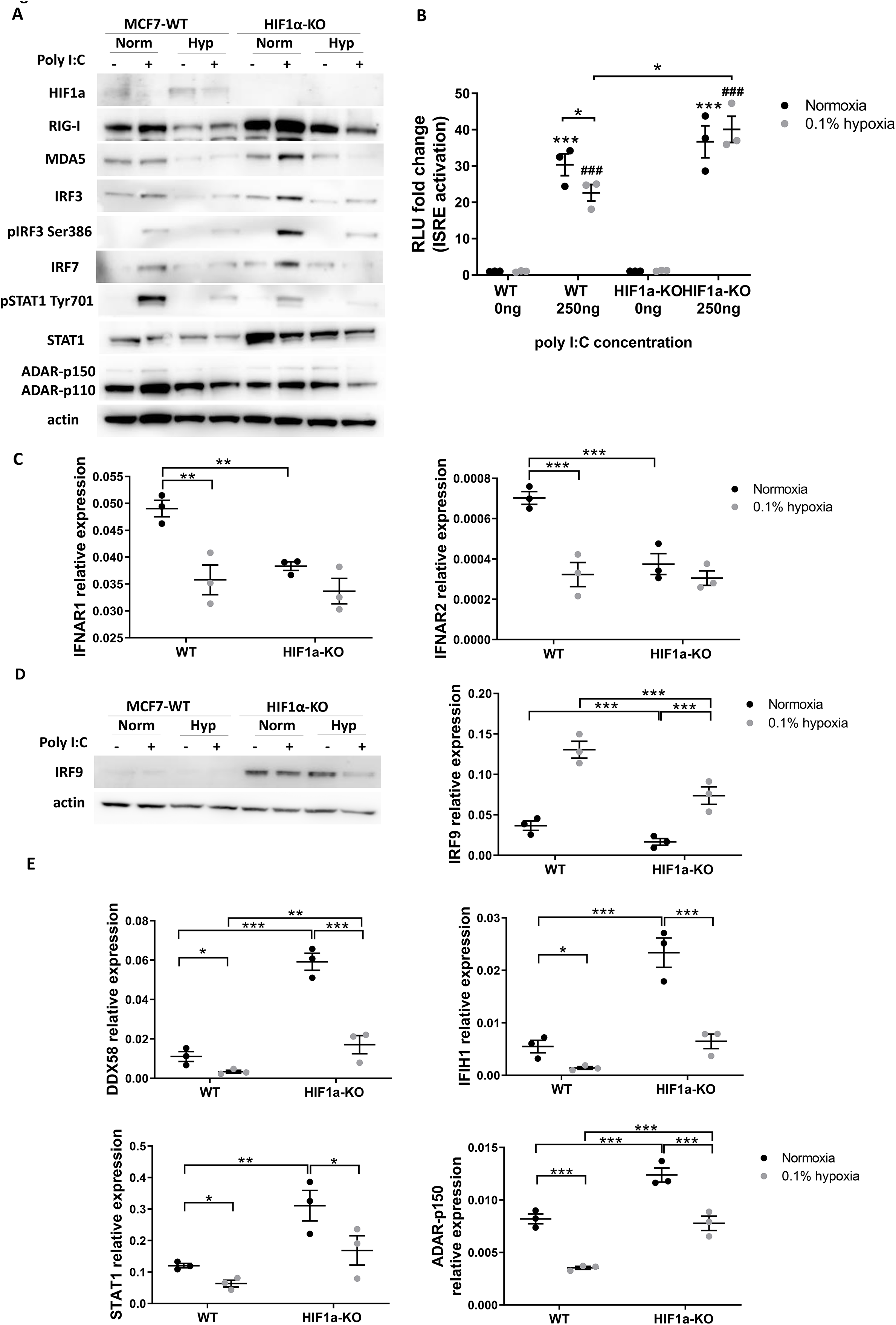
HIF1α-independency of type I IFN downregulation in hypoxia. A) Western-blot showing protein changes in different MCF7-WT vs MCF7-HIF1α-KO ells when exposed to normoxia or 0.1% hypoxia for 48h and transfected with 0.5ug/ml poly I:C in the last 6h of the treatment B) ISRE activation via luciferase ctivity in 3C11 cells using supernatants of MCF7-WT and MCF7-HIF1α-KO cells previously treated with normoxia or 0.1% hypoxia for 24h and transfected with poly C (n=3). C) IFNAR1 and IFNAR2 mRNA expression in MCF7-WT and MCF7-HIF1α-KO cells. D) IRF9 protein and mRNA levels in MCF7-WT and MCF7-HIF1α-O cells in the same as experiment as in A). E) mRNA expression in MCF7-WT and MCF7-HIF1α-KO cells cultured in normoxia or 0.1% hypoxia for 48h. * p<0.05, * p<0.01, *** p<0.001 in normoxia and normoxia vs hypoxia, # p<0.05, ## p<0.01, ### p<0.001 in hypoxia samples.

The effect of poly I:C in hypoxia was attenuated for all the above proteins to similar residual levels in both cell lines, which excluded HIF1α as the main mechanism.

Higher IRF3, pIRF3 and IRF7 induction in MCF7-HIF1α-KO than in MCF7-WT suggested a possible increase in IFN production. Using the supernatant from these cells to treat 3C11 cells as described above, there was no ISRE activation in normoxic supernatants from WT or MCF7-HIF1α-KO at basal level. For both, ISRE activation was similarly increased by poly I:C in normoxia. However hypoxic MCF7-WT supernatant caused a significantly lower ISRE stimulation and this decrease was not noted for hypoxic MCF7-HIF1α-KO supernatants which triggered significantly higher ISRE activation than MCF7-WT (figure 5B).

*IFNAR1* and *IFNAR2* expression was determined and they were significantly downregulated in MCF7-WT cell in hypoxia compared to normoxia. Interestingly, MCF7-HIF1α-KO cells showed in normoxia significantly lower expression of both compared with MCF7-WT and comparable to hypoxic MCF7-WT levels. Moreover, hypoxia did not further decrease *IFNAR1* and *IFNAR2* levels at mRNA level (figure 5C). This shows that HIF1α is unlikely to explain the downregulation in hypoxia, as *IFNAR1* and *IFNAR2* were lower without HIF1α, and they would be expected to be higher.

These data suggest a complex mechanism in which HIF1α has a basal role increasing STAT1 and responsiveness to poly I:C, but the effect of hypoxia downregulating many IFN genes persists. Finally, the hypoxia downregulation of IFN secretion is lost, implying a role for HIF1α in this last step.

Some ISGs associated with chronic anti-viral response such MDA5, MX1 or STAT1 can be transcribed by unphosphorylated STAT1, and this effect is driven by higher IRF9 expression (29). Both at mRNA and protein levels, MCF7-HIF1α-KO cells showed higher IRF9 expression (figure 5D) associated with significantly higher ISG expression even in hypoxia compared to MCF7-WT cells (figure 5E).

However, as for all the other cell lines, hypoxic MCF7-HIF1α-KO cells showed lower ISG expression than normoxic MCF7-HIF1α-KO cells. These data suggest that HIF1α upregulation in hypoxia is partly responsible for the type I IFN pathway downregulation observed in this condition but other HIF1α-independent mechanisms cause a further decrease.

### Type I IFN downregulation and role of HIF2α

The converse experiment was performed with renal cancer cells, which showed vHL loss or mutations leading to stabilization of HIF1α/HIF2α in normoxia. RCC4-EV (empty vector, mutant vHL), RCC4-vHL (transfected with wild type vHL and, as a result, displaying lower levels of HIF1α/HIF2α in normoxia), 786-0 parental (vHL mutation and HIF1α deficient) and 786-0-HIF2α-KO (deficient in both HIF1α and HIF2α) were used. In RCC4 cells, poly I:C strongly induced MDA5 in normoxia and this was reduced in hypoxia, showing that high HIF1α/HIF2α alone was not enough to suppress normoxic levels to those obtained in hypoxia. Poly I:C induced similar changes in RIG-I, IRF3 and pIRF3 in both cell lines in normoxia, again showing no effect of high HIF1α/HIF2α in this circumstance. Much higher STAT1 and pSTAT1 was detected in the vHL corrected cell line in normoxia and both decreased in hypoxia, clearly linking suppression of STAT1 to HIF1α/2α (Supplementary figure 3A). In this model pSTAT1 was upregulated and STAT1 downregulated in RCC4-wild type vHL cells, also pointing to a role of HIF1α/2α in suppressing phosphorylation.

In 786-0 cells (Supplementary figure 3B), poly I:C did not affect RIG-I or MDA5 levels but it induced pIRF3 and pSTAT1 expression in both cell lines. As observed in other HIF1α-KO cell lines, STAT1 level was highly expressed in both cell lines. However, hypoxia led to downregulation of MDA5, IRF3, pIRF3, STAT1, pSTAT1 and ADAR expression, again showing that constant deficiency of HIF1α/HIF2α still allowed a further suppression of IFN response in hypoxia.

In general and despite of the differences observed in these models, there is a partial role of HIF1/2α in regulating some members of the pathway, but more importantly, it is clear that there is a HIF1/2α-independent downregulation of the type I IFN pathway.

### Type I IFN pathway is also downregulated in hypoxic areas *in vivo*

To confirm that hypoxic areas *in vivo* also showed lower expression of type I IFN pathway, xenograft experiments were conducted using MCF7 and MDA-MB231 cells. Avastin, a VEGF antagonist, was used to create tumour hypoxia by reducing angiogenesis, as described previously (30–32). Immunohistochemistry was performed using antibodies against IRF3, IRF7 and ADAR in 5 control and 5 avastin-treated mice for each cell line. Avastin treatment decreased the percentage of CD31-positive cells (vessel marker) and, consequently, the proportion of necrosis was significantly higher (25). Moreover, the percentage of CA9+ cells was greater in Avastin-treated xenografts compared to PBS. Interestingly, IRF3, IRF7 and ADAR were significantly downregulated in avastin-treated mice in the case of MCF7 xenografts (figure 6A), and IRF3 and IRF7 also showed significantly lower expression in MDA-MB231 xenografts (figure 6B).

**Figure 6.**
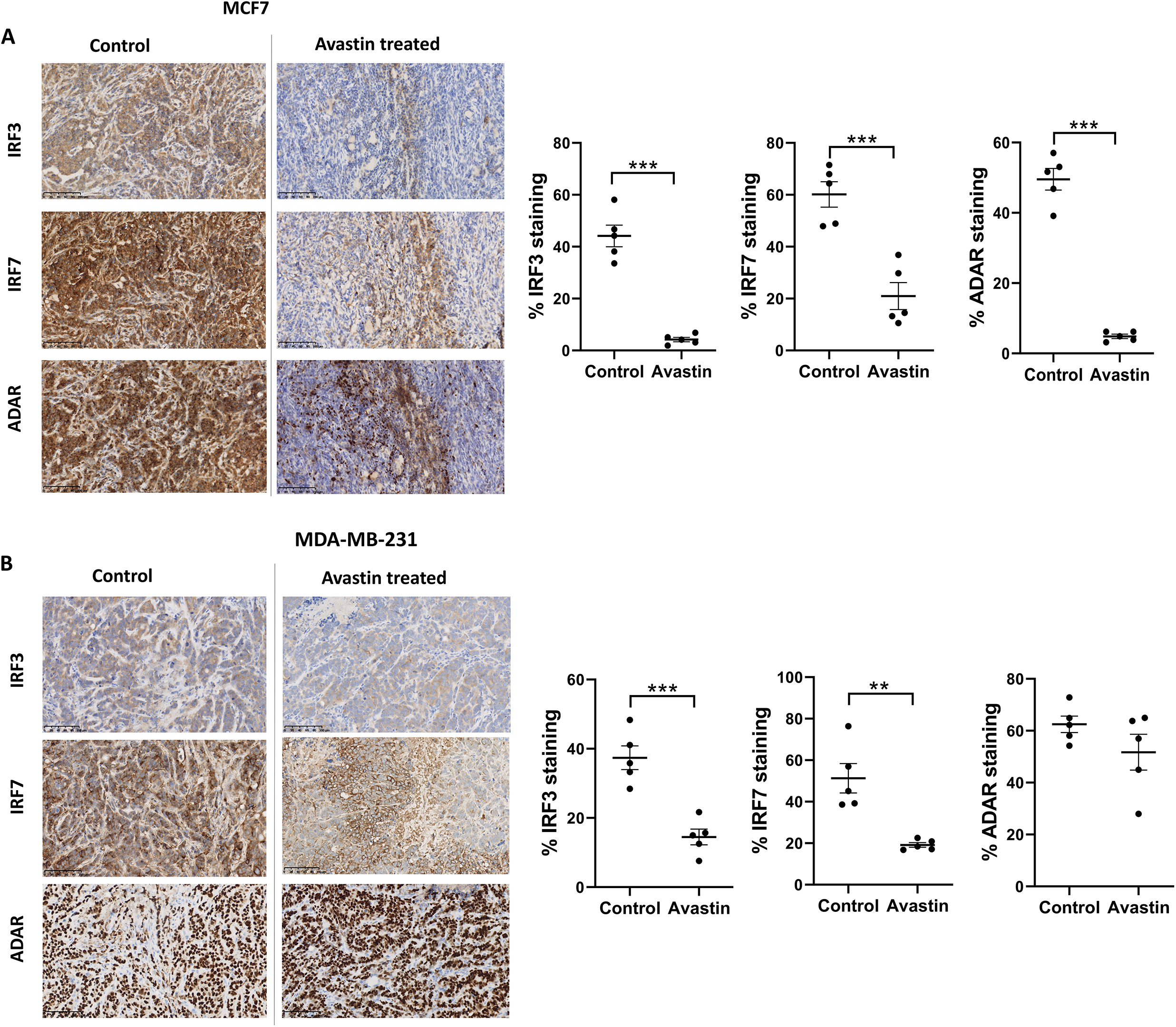
Downregulation of the type I IFN staining in hypoxic tumours *in vivo*. A) IRF3, IRF7 and ADAR staining in MCF7 xenografts from control and avastin-treated mice and their respective quantification. B) IRF3, IRF7 and ADAR staining in MDA-MB-231 xenografts from control and avastin-treated mice and their respective quantification. n=5 per group, ** p<0.01, *** p<0.001.

### ATAC-Seq data shows lower chromatin accessibility on STAT1 and IRF3 containing promoters

To investigate how hypoxia may downregulate the type I IFN pathway independently of HIF1α/HIF2α we performed ATAC-seq on MCF7 cells cultured for 48h in normoxia or 0.1% hypoxia. We found 5,577 peaks (7.4%) with differential accessibility (figure 7A). Peaks showing increased accessibility during hypoxia (n = 2,439) were enriched for promoters, whereas those with decreased accessibility (n = 3,138) tended to be intergenic or intronic (figure 7B), showing a switch in regulatory networks driving gene expression. Consistent with HIF1/HIF2 driving an active hypoxia response, motif analysis showed enrichment for HIF1 sites in distal peaks with increased accessibility (figure 7C). Conversely, peaks with differential decreased accessibility showed higher levels of STAT1, IRF3, FOXA1 and GATA3 motifs, than unchanged peaks. FOXA1 and GATA3 motifs were present more frequently in distal peaks with decreased accessibility and less frequently present in peaks with differential increase accessibility. IRF3 motifs were significantly less represented in peaks with increased accessibility both at promoters and distal sites. STAT1 motifs only showed significantly less presence in increased accessible peaks at promoters. The observed loss of accessibility at FOXA1 and GATA3 sites is consistent with their roles downstream in ER signaling and the degradation of ER under low O2 conditions (33). Therefore, hypoxia appears to drive global changes in chromatin accessibility, partially through the shutdown of both type I IFN and ER signalling responsive promoters and enhancers, although globally most decreases are in intergenic regions and introns.

**Figure 7.**
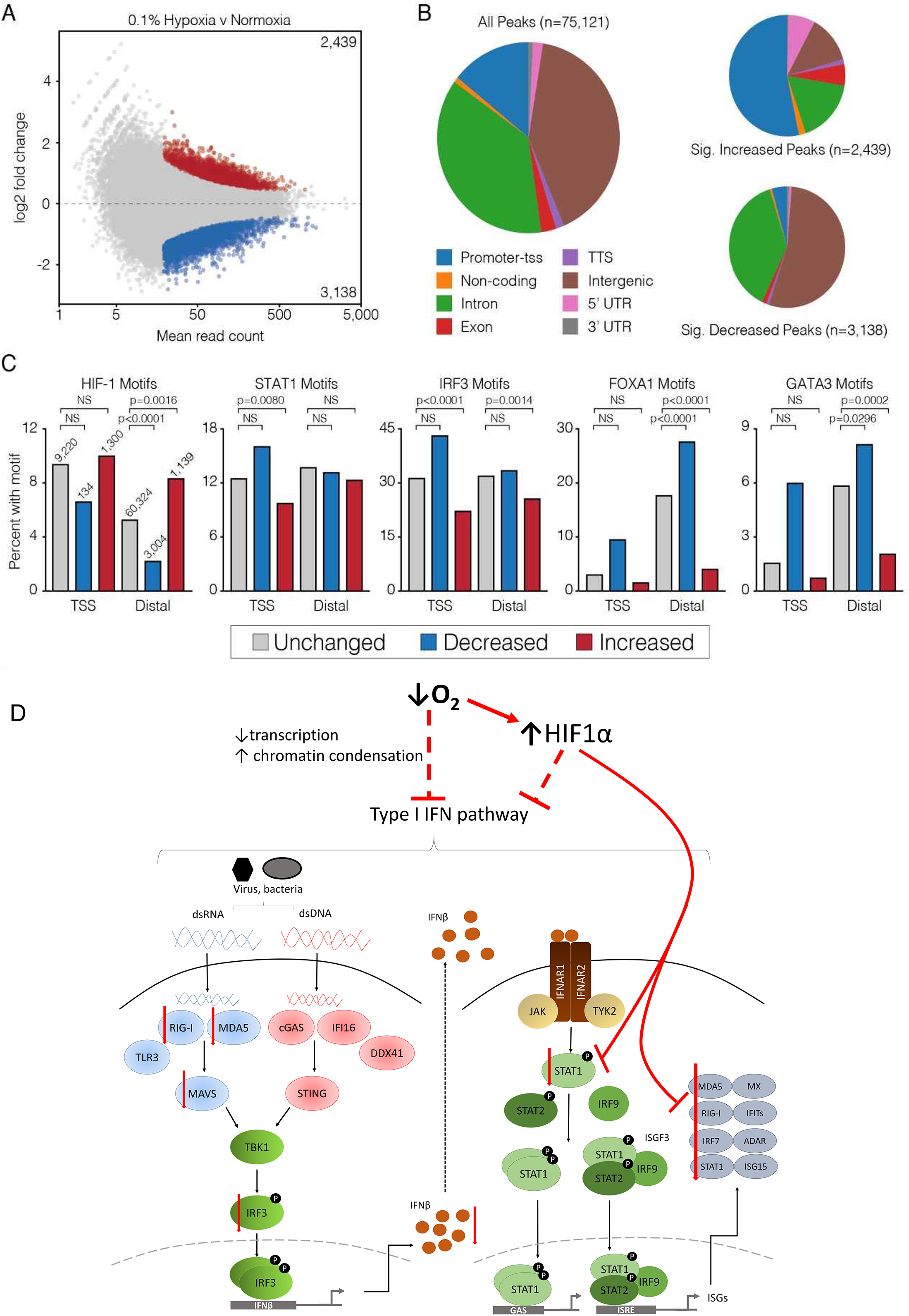
ATAC-seq reveals dynamics of chromatin accessibility during hypoxia. A) MA plot of differentially accessible peaks during hypoxia with peaks significantly more (red) or less (blue) accessible during hypoxia. B) Annotation of all ATAC-seq peaks, and peaks with significant changes in accessibility. C) Percent of ATAC-seq peaks containing HIF1, STAT1, IRF3, FOXA1 and GATA3 transcription factor binding motifs. Peaks are divided into transcription start site (TSS) associated and TSS-distal peaks (all other categories in B). The number of peaks that are unchanged, decreased and increased in A) are shown graphically and following the same colour-code in C). Number of differentially accessible peaks in each class is shown in the HIF-1 graph and are identical for the others. P-values are for a Fisher’s exact test with Bonferroni multiple test correction. D) Schematic representation of the hypoxia-dependent and HIF1α-independent mechanism by which hypoxia downregulates the type I IFN pathway.

## DISCUSSION

We describe a new mechanism of hypoxic immunosuppression via downregulation of the type I IFN pathway. This occurred at oxygen tensions commonly found in tumours and in more severe hypoxia present in many cases (1% vs 0.1%). The type I IFN pathway is downregulated at RNA level in less than 16h under hypoxia whereas protein level is downregulated after a 48h exposure. Reoxygenation gradually reverses the levels and it takes 24h to reach normoxic levels again. The observation that RNA recovers more quickly than protein after reoxygenation points to *de novo* transcription or stabilisation occurring during the reoxygenation phase, and RNA changes leading to the protein expression changes rather than protein degradation. However, there was no overshoot and upregulation above the normoxic baseline after reoxygenation.

Furthermore, we observed lower activation of the type I IFN pathway and less production/secretion of interferons under hypoxia in all cell lines tested when they were activated with poly I:C. This suggests that endogenous activators of the pathway in a hypoxic tumour microenvironment would lead to lower stimulation and consequently to lower immune response. This hypothesis is supported by lower presence of T-cells and NK cells in hypoxic areas in lung and dermal tumours (34) and in prostatic tumours (35). In both studies the immune infiltration increased when the hypoxic areas were reduced using respiratory hyperoxia (34) or the hypoxia-activated prodrug TH-302 (35), respectively.

We observed that the type I IFN downregulation in hypoxia is independent of HIF1α and HIF2α, although some members (RIG-I, and mainly STAT1) are regulated by HIF1α as observed in MCF7-HIF1α-KO cells. Moreover, HIF2α seems not to play an important role in the type I IFN downregulation as observed in 786-0 HIF2α-KO cells. Previous work reported that RIG-I harbours hypoxic response elements in its 3’UTR and RIG-I expression was upregulated during hypoxia in human myotubes (36). However, a different study in tumour cells supported our finding that RIG-I and MDA5 are downregulated under hypoxia and they required functional IFNAR1 to respond to stimulus (37). Recently, type I IFN signalling was shown to be inhibited by high lactate levels generated during the metabolic switch to glycolysis occurring in hypoxia (38). As lactate production requires HIF1α, this result could explain the partial effect we observed in HIF1α-KO cells. However, the inhibitory effect seen in hypoxia needed 16h to be significant and it could be a consequence of the HIF1α-independent general transcription inhibition occurring during hypoxia that leads to 42% and 65% decrease in total RNA transcription after 24h and 48h exposure to 0.2% hypoxia, respectively (39).

Apart from the transcriptional inhibition, translation blockade also occurs in hypoxia as a quick and reversible mechanism upon reoxygenation via phosphorylation of eIF2α to adapt to changing environmental conditions. This mechanism is also HIF1α-independent and peaks after 2h in hypoxia but often recovers after 24 hours (40, 41).

We investigated each step in the type I IFN pathway, from the sensors (RIG-I, MDA5), transcription factors (IRF3, IRF7, STAT1), ISGs (MX1, ADAR) and interferon release, and each stage was reduced under hypoxic conditions, although there was heterogeneity. We therefore studied the chromatin accessibility in hypoxia related to these genes by ATAC-Seq. Overall we found that 48h exposure to hypoxia led to global changes rather than changes in specific pathways and only 7% of the type I IFN pathway were associated with differentially accessible peaks. Interestingly, hypoxia caused decreased accessibility at STAT1 and IRF3 containing promoters, suggesting lower transcription from those genes but the downregulation and lower activation observed in hypoxia cannot be explained only by this. However, a recent paper showed that 1h exposure to hypoxia caused a rapid and HIF1α-independent induction of histone methylation, and the locations of H3K4me3 and H3K36me3 predict the transcriptional response hours later (42). These methylation markers are normally related to active transcription and they were significantly less present in type I IFN genes after hypoxia, potentially indicating that the expression of genes in this pathway are repressed by histone modifications that occurred in hypoxia due to decreased enzyme activity of oxygen-requiring JmjC-containing enzymes, and particularly KDM5A, that would affect transcription and translation of this pathway later on (42).

Here, we propose a model in which hypoxia downregulates every single step in the type I IFN pathway. However, there is a partial effect of HIF1α as its deletion increased the levels of MDA5, RIG-I, IRF3, IRF7, STAT1 and IFN secretion upon activation in normoxia by upregulating IRF9 and not by increasing phosphorylation of STAT1. Conversely, hypoxia is still able to decrease the IFN response when HIF1α is deleted potentially by specifically decreasing the transcription of this pathway (figure 7D).

Supporting our work, ISG15 (a typical ISG increased by IFNαβ) was reported to decrease in hypoxia and interact and modify HIF1α via ISGylation, thus reducing its levels and affecting its biological impact on EMT and cancer stemness acquisition (43). In the same line, response to type I IFN pathway was reported to be incompatible with induction of pluripotency (44).

All together, these data suggest that the relevance of this hypoxic downregulation of the IFN response is the potential effect on enhancing tumour growth by decreasing the cytotoxicity of T lymphocytes and the capacity of dendritic cells (DCs) to process and present antigens (11). As a result, this downregulation could contribute to the radio- and chemo-resistance observed in hypoxic areas as these therapies rely on intact type I IFN signalling (14). This provides a further rational for targeting the hypoxic tumour population in combination with check point inhibitor therapy (45) and selection of hypoxic patients for study of IFN response inducers is warranted.

## ACKNOWLEDGEMENTS

We thank Dr Dimitrios Anastasiou and Dr Fiona Grimm (The Francis Crick Institute, UK) for giving us MCF7-WT and MCF7-HIF1α-KO cells.

We also thank Matthew Gosden (University of Oxford, UK) for his technical advice performing the ATAC-Seq experiments.

## AUTHOR CONTRIBUTION

AM and ALH designed the experiments, analysed the data and wrote the manuscript. AM and EA performed the experiments. EB performed the animal experiments and stained the samples. SB performed the single cell experiments and APC analysed them. DD and RB analysed ATAC-Seq data, and JR provided reagents and advice. This research was supported by funding from Cancer Research UK (ALH, EB, SB), Breast Cancer Research Foundation (ALH), and Breast Cancer Now (ALH, AM).

