## supplementary information for "Hypoxia induces transcriptional and translational downregulation of the type I interferon (IFN) pathway in multiple cancer cell types"

Supplementary table 1. Cell lines used.

| Cell line | Source | Reference |
|---|---|---|
| MCF7 | ATCC | Cat# HTB-22, RRID:CVCL_0031 |
| T47D | ATCC | Cat# HTB-133, RRID:CVCL_0553 |
| BT474 | ATCC | Cat# HTB-20, RRID:CVCL_0179 |
| MDA-MB-231 | ATCC | Cat# CRM-HTB-26, RRID:CVCL_0062 |
| MDA-MB-453 | ATCC | Cat# HTB-131, RRID:CVCL_0418 |
| MDA-MB-468 | ATCC | Cat# HTB-132, RRID:CVCL_0419 |
| HCC1187 | ATCC | Cat# CRL-2322, RRID:CVCL_1247 |
| RCC4 | ECACC | Cat# 03112702, RRID:CVCL_0498 |

Supplementary table 2. Antibodies.

| Protein | Antibody | Company | Dilution |
|---|---|---|---|
| actin | | Sigma | 1:50000 WB |
| ADAR | #14175 | Cell Signaling | 1:1000 WB<br>1:200 IHC |
| HIF1a | 610958 | BD Transduction Lab | 1:1000 WB |
| HIF2a | NB100-122 | Novus | 1:500 WB |
| IRF3 | Ab76409 | Abcam | 1:1000 WB<br>1:200 IHC |
| IRF3 (phospho Ser386) | Ab76493 | Abcam | 1:1000 WB |
| IRF7 | Ab109255 | Abcam | 1:1000 WB<br>1:200 IHC |
| IRF9 | #76684 | Cell Signaling | 1:1000 WB |
| MAVS | ALX-210-929-C100 | Enzo Life Sciences | 1:1000 WB |
| MDA5 | In house | | 1:500 WB |
| STAT1 | #9172 | Cell Signaling | 1:1000 WB |
| STAT1 (phospho Tyr701) | #9167 | Cell Signaling | 1:1000 WB |
| STAT1 (phospho Ser727) | #9177 | Cell Signaling | 1:1000 WB |
| RIG-I | LS-C331000 | LS Bio | 1:1000 WB |

Supplementary table 3. Primers.

| Primer | Sequence |
|---|---|
| ADAR_p150_F | CTTCCAGTGCGGAGTAGCG |
| ADAR_p150_R | GTGACGGTGTCTGCTTTCCA |
| DDX58_F | CAAGCCTTCCAGGATTATATCCG |
| DDX58_R | AGTCCAGAATAACCTGCATGGT |
| IFIH1_F | TTAACAGGCTCTGATTGCTC |
| IFIH1_R | TCTCTTCATCTGAATCACTTCCC |
| IFNAR1_F | ATTTCGCAAAGCTCAGATTGGT |
| IFNAR1_R | CATCCAAAGCCCACATAACAC |
| IFNAR2_F | GCCAGGCCTCAGAATCAGCA |
| IFNAR2_R | TGTAAATGACCTCCACCATATCCA |
| IRF3_F | CTCGTGATGGTCAAGGTTGTG |
| IRF3_R | AGTTTATTGGTTGAGGTGGTGG |
| IRF7_F | AAGGGCTTCCCCTGACTG |
| IRF7_R | TCTACTGCCCACCCGTACA |
| IRF9_F | AGAAAGTACCATCAAAGCGAC |
| IRF9_R | TTGTGTCTGTAACTTCCTGTG |
| MAVS_F | GTACCCGAGTCTCGTTTCCT |
| MAVS_R | ATGAAGTACTCCACCCAGCC |
| MX1_F | GTTACCAGGACTACGAGATTGAG |
| MX1_R | GATGAGTGTCTTGATCTTATACCC |
| PKR_F | ATCTGACTACCTGTCCTCTG |
| PKR_R | GAGACCATTCATAAGCAACGA |
| RPL11_F | GCAAACTCTGTTCAACATCTG |
| RPL11_R | CATACTCCCGCACTTTAGAC |
| STAT1_F | TACACCTACGAACATGACCCT |
| STAT1_R | TCACCAACAGTCTCAACTTCAC |
| STAT2_F | CCCGCTGACTGAAATCATCC |
| STAT2_R | AGTTCATCCACCTGTCTATTAGAG |

Western blot

Cells were washed with cold PBS and lysed 30 min on ice with RIPA lysis buffer (R0278, Sigma) containing protease (cOmplete, 11697498001) and phosphatase (phosSTOP, 4906845001) inhibitor cocktails (Roche). Lysates were cleared by centrifugation and supernatants were boiled at 95°C for 5min in 4x NuPAGE LDS sample buffer (NP0007, Invitrogen) containing 10% β-mercaptoethanol. Samples were run on NuPAGE Novex 4-12% Bis-TRIS gels (NP0336BOX, Invitrogen) using NuPAGE MOPS-SDS running buffer (NP000102, Invitrogen). After transferring the proteins onto PVDF membranes (IPVH00010, Millipore), these were blocked with 5% skimmed milk (A0830,100,

Panreac) in TBS-T (TBS/0.1% Tween-20) for 1 hour at room temperature and were then incubated overnight with primary antibodies (supplementary table 2) in 3% milk TBS-T at 4°C. Membranes were washed 3X in TBS-T and incubated with HRP-anti-mouse/rabbit secondary antibodies (Gibco). Development was performed with Amersham ECL Prime Western Blotting Detection Reagent (GERPN2232, GE Healthcare Life Sciences) using ImageQuant™ LAS 4000. Stripping with Restore PLUS Western Blot Stripping Buffer (46430, Themor Fisher Scientific) was performed to blot different antibodies in the same membrane.

RT-qPCR

RNA from each experiment was extracted using the Tri-Reagent protocol (T9424, Sigma) and 1µg of RNA was reverse transcribed into cDNA with the High Capacity cDNA reverse transcription kit (44368813, Thermo Fisher Scientific) using random hexamer primers. The PCR reaction containing SensiMix™ SYBR Green® No-ROX Kit (QT650-20, Bioline) was run on a 7900 Real time PCR System (Applied Biosystems) with standard cycling conditions: 10 minutes 95°C, and 40 cycles of 15 seconds 95°C followed by 1 minute 60°C. Gene expression was analysed with the Ct method using *RPL11* expression for normalization. The primers used are listed in Supplementary table 3.

IFN bioassay

HEK293-3C11 cells stably transduced with an ISRE-Luc reporter construct (1) were used to detect possible IFN present in the supernatant. $2x10^4$ MCF7 cells/well were transfected in triplicate with different concentrations of poly I:C and washed with PBS after 1h, cells were incubated for 24h in normoxia or 0.1% hypoxia and 24h after, the supernatant was transferred into $2x10^4$ 3C11 cells kept in normoxia. After 24h incubation, 3C11 cells were lysed and measured using OneGlo luciferase assay (E6120, Promega) in a FluorOPTIMA luminometer.

Single Cell sequencing and analysis

Single-cell RNA-seq libraries were prepared as per the Smart-seq2 protocol by Picelli et al (2) with minor technical adaptations detailed in Suppementary material. Reverse transcription was carried on with 0.25 U/reaction of SuperScript II (Invitrogen) and PCR pre-amplification with 5′ biotinylated IS PCR primers (Biomers) for 22 cycles. PCR cleaning was performed by using Ampure XP Beads (Beckman Coulter) at a ratio 0.8:1 (beads:cDNA). cDNA was re-suspended in elution buffer (Qiagen) and assessed with a High-sensitivity DNA chip (Agilent). Barcoded Illumina sequencing libraries (Nextera XT, Library preparation kit, Illumina) were generated using the automated platform (Biomek FXp). Finally, libraries were pooled, and sequencing was performed in paired-end mode for 2×75 bp cycles using the Illumina HiSeq 4000 sequencer.

Single cell sequencing analysis

scRNA-seq data were deposited in Gene Expression Omnibus under SuperSeries accession number GSE134038. Following deconvolution of RNA-seq raw reads into individual cells, initial quality assessment was performed using *FastQC* (v0.11.4). Reads pairs were mapped to a composite genome made by concatenating the human genome reference sequence GRCh37 (hg19) and 92 ERCC ExFold RNA Spike-In Mixes sequences (ThermoFisher). Alignment was performed using *HISAT2* (v2.0.3b) in default settings. Uniquely aligned read pairs were assigned and counted towards annotated human genomic genes and ERCC RNA spike-in features using *featureCounts* (v1.5.0-p2). All metrics were subsequently collected using *MultiQC* (v0.8) for reporting. Filtering was performed with Seurat v2.1 and manually by selecting on median absolute deviations (MADs) thresholds. 66 cells (17%) were identified as outliers and removed from all downstream analyses.

Differentially expressed gene analysis was performed using the nonparametric Wilcoxon test on $\log_2$(TPM) expression values for the comparison of expression level and Fisher′s exact test for the comparison of expressing cell frequency. *P* values generated from both tests were then combined using Fisher's method and were adjusted using Benjamini–Hochberg (BH). Differentially expressed genes were selected on the basis of the absolute $\log_2$ fold change of ≥1 and the adjusted *P* value of <0.05. Selected genes were subjected to the hierarchical clustering analysis using Pearson correlation as a distance, with heatmap generation performed using the *pheatmap* package in R.

Xenograft growth and IHC

Procedures were carried out under a Home Office license. Xenograft experiments were performed as described in (3). A standard haematoxylin and eosin protocol was followed to assess the morphology and the amount of necrosis on xenografts. For immunohistochemistry staining, slides were dewaxed and antigen retrieval performed in pH 6. Slides were stained using the FLEX staining kit (Agilent). To summarise, endogenous peroxidase activity was blocked before slides were stained with rabbit antibodies diluted as indicated in supplementary table 2 for 1h at room temperature. Slides were incubated with the Flex anti-rabbit/anti-mouse secondary antibody for 30 minutes at room temperature and washed in Flex buffer. 3,3′-Diaminobenzidine (Flex-DAB) was applied to the sections for 10 minutes. The slides were counterstained by immersing in Flex-hematoxylin solution for 5min, washed and air-dried before mounted with mounting medium (Sigma). Secondary-only control staining was done routinely, these were negative.

Expression of markers and viable/necrotic areas was quantified on whole sections quantitatively by using the Visiopharm Integrator System. HDAB-DAB colour deconvolution band is used to detect positively stained cells. Threshold classification is used to identified necrosis and living regions and thus identify number of positive staining within these regions. Appropriate thresholds levels are checked against control xenografts staining before being set and the xenografts from all groups are then analysed.

ATAC-seq

MCF7 cells exposed to normoxia or 0.1% hypoxia for 48h were used. 75000 cells per technical replicate (x2) in 3 biological replicates were lysed in 50ul cold lysis buffer (10mM Tris-HCl pH 7.5, 10mM NaCl, 2mM $MgCl_2$, 0.1% Igepal CA-630). Nuclear pellets were obtained after 10 min centrifugation at 4°C at 500G and resuspended in 50ul of tagmentation mix (25ul 2x TD Buffer, 2.5ul Tn5 transpose and 22.5ul nuclease free $H_2O$) with the Nextera Kit (FC-121-1030, Illumina), then incubated for 30 min at 37°C. DNA was purified using the Qiagen MinElute columns (28004, Qiagen). The samples were indexed using 10ul of transposed DNA, 10ul nuclease free $H_2O$, 2.5ul 25uM customised Nextera PCR primer 1, 2.5ul 25uM customised nextera primer 2 (barcode in supplementary table 3) and 25ul NEB Next High-Fidelity 2x PCR Master Mix (M0541S, NEB). The PCR was performed with the following protocol: 72°C for 5 min, 98°C for 30 sec, 11 cycles of 98°C for 10 sec, 63°C for 10 sec and 72°C for 1 min. The PCR product was purified with Qiagen PCR Cleanup Kit (28104, Qiagen). The samples were sequenced on a next generation sequencing platform using the NextSeq® 500/550 High Output Kit v2 (75 cycles; FC-404-2005, Illumina). ATAC-seq data is available in the Gene Expression Omnibus (GSE133327).

ATAC-Seq and ChIP-Seq analysis

ChIP-seq for HIF-1α, HIF-1β, HIF-2α (GSE28352 (4)) and FOXA1 (GSE80808, GSE25710 (4, 5)) were downloaded from the sequence repository archive (https://www.ncbi.nlm.nih.gov/sra). ChIP-seq and ATAC-seq reads were mapped (hg19) and filtered for PCR duplicates with NGseqBasic (Telenius et al, http://dx.doi.org/10.1101/393413*)* which uses bowtie and samtools. Regions of ATAC-seq open chromatin were identified and filtered to exclude chrM, chrX, chrY and large copy number variations. Read coverage was calculated using deepTools (6). Open chromatin regions were annotated using HOMER (7). Changes in chromatin accessibility were determined using DESeq2 and differential peaks were identified using an adjusted p-value threshold of 0.05. Transcription factor motifs were downloaded from HOCOMOCO (8) and identified in ATAC-seq peaks using FIMO (9). Background motif frequency was identified from ATAC-seq peaks with a padj>0.3, absolute log2FoldChange<0.25 and baseMean>30. Motif enrichment in differential peaks relative to background was calculated using Fisher's exact test with Bonferroni correction.

**Supplementary Figure 1**

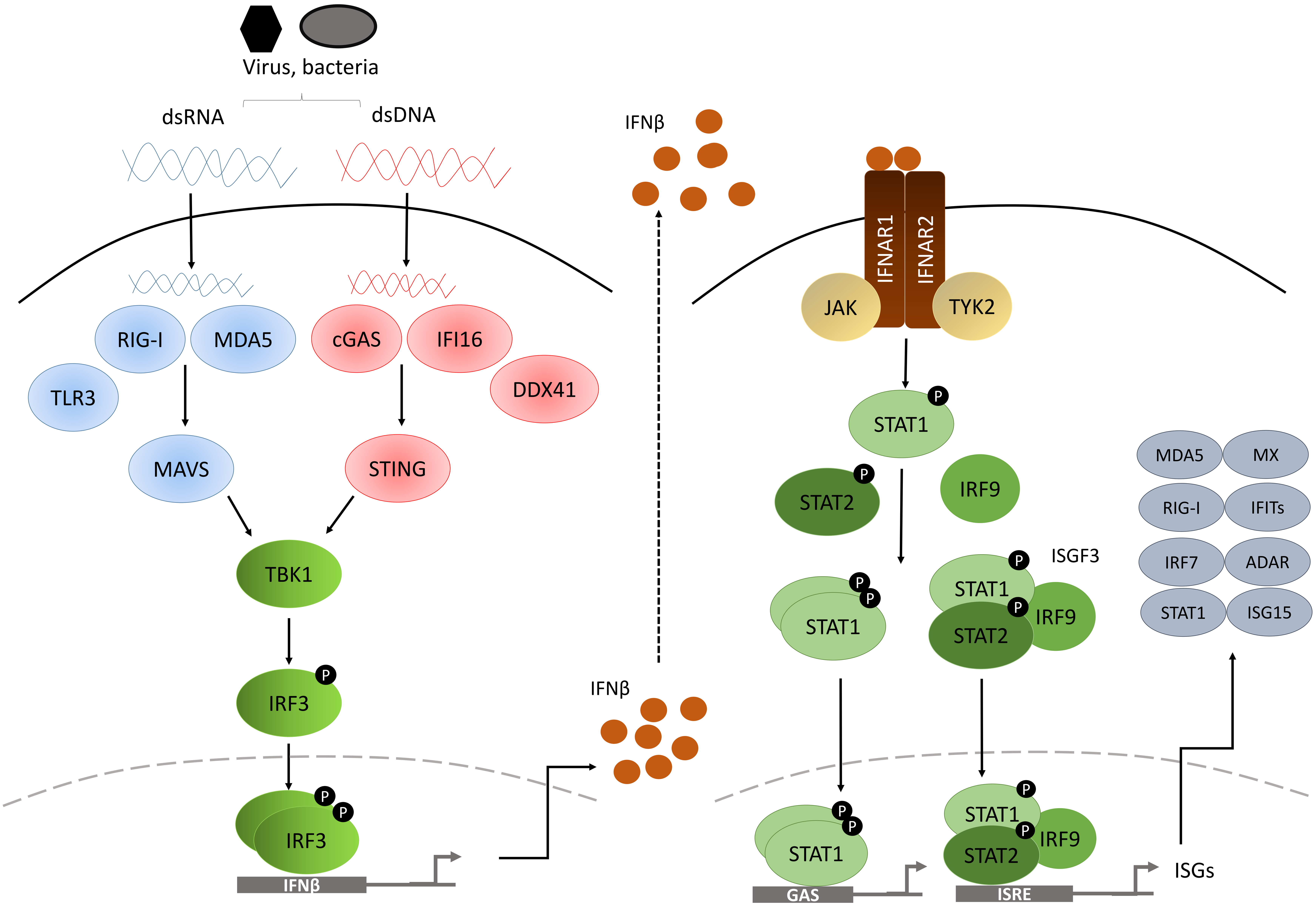


**Supplementary figure 1. Type I IFN signalling.** Type I IFN pathway senses bacterial or viral double-stranded DNA (dsDNA) or unmethylated DNA (through cGAS, DDX41, IFI16) and dsRNA (through RIG-I, MDA5, TLRs) in the cytoplasm. The adaptors MAVS (for RNA) and STING (for DNA) trigger the pathway activating TBK1 which phosphorylates IRF3. Phosphorylated IRF3 forms dimers which translocate into the nucleus to activate the transcription of IFNα/β promoter. IFNs are secreted and bind to its receptor (IFNAR1/2) in the plasmatic membrane in an autocrine or paracrine manner. Then, the JAK-STAT1 pathway is triggered leading to the complex ISGF3 complex (STAT1-STAT2-IRF9) which translocates into the nucleus to activate the transcription of INF-stimulated genes (ISGs).

**Supplementary Figure 2**

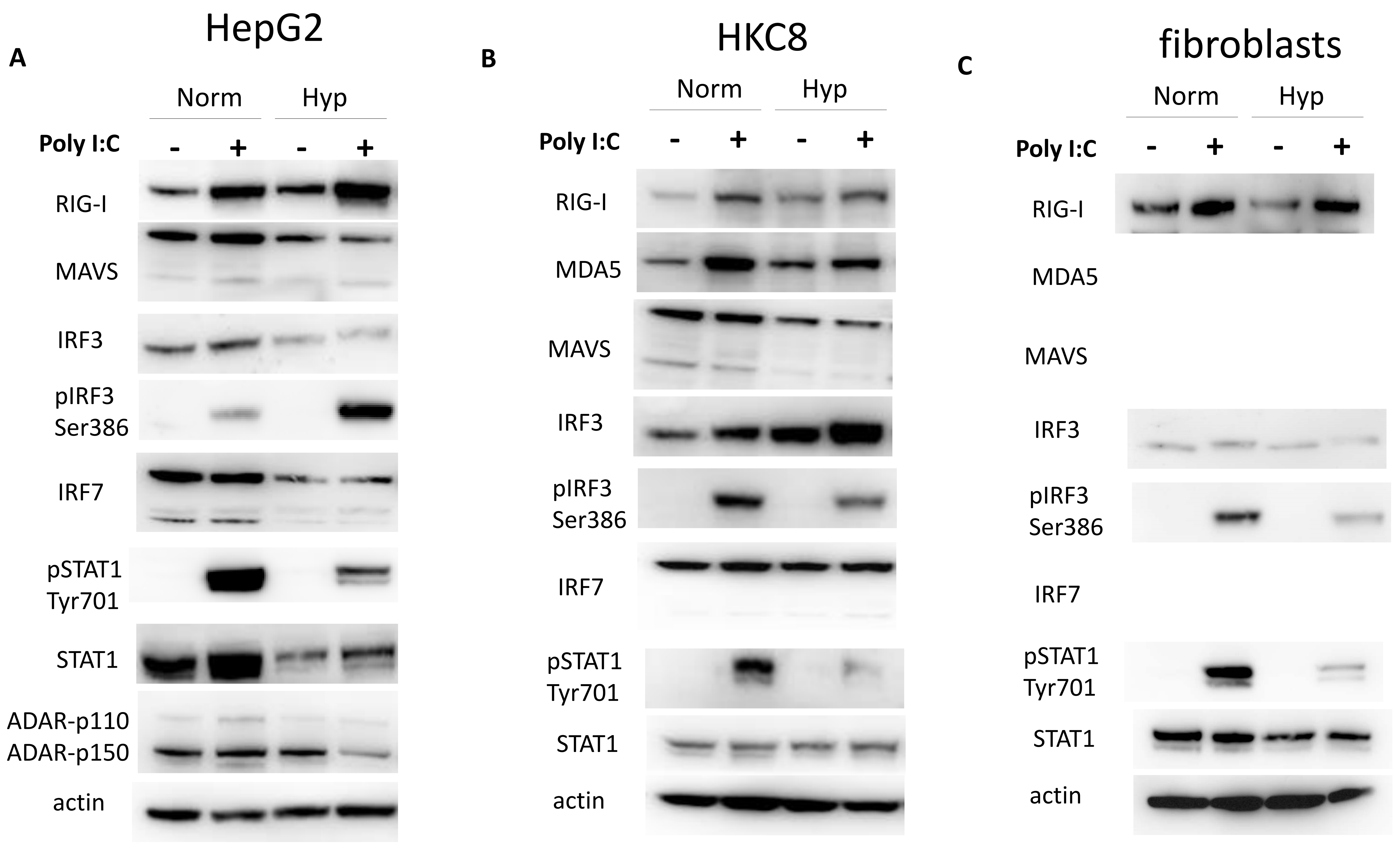


**Supplementary figure 2. Type I IFN expression in other cancer models**. A) Protein levels of type I IFN pathway members in HepG2, B) HCK8 cells, and C) fibroblasts after being exposed to normoxia or 0.1% hypoxia for 48h and transfected with poly I:C in the last 6h (n=3).

**Supplementary figure 3**

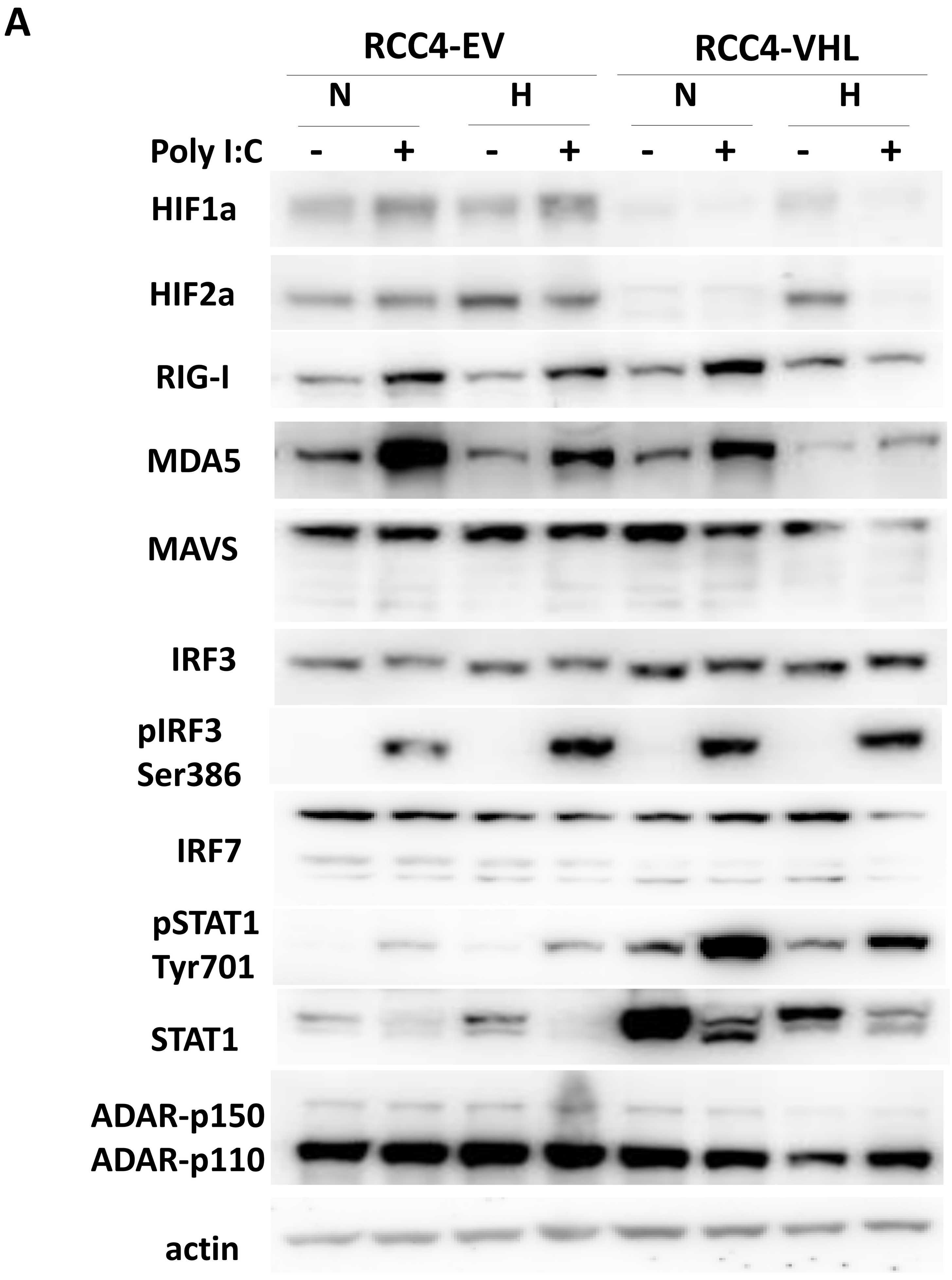


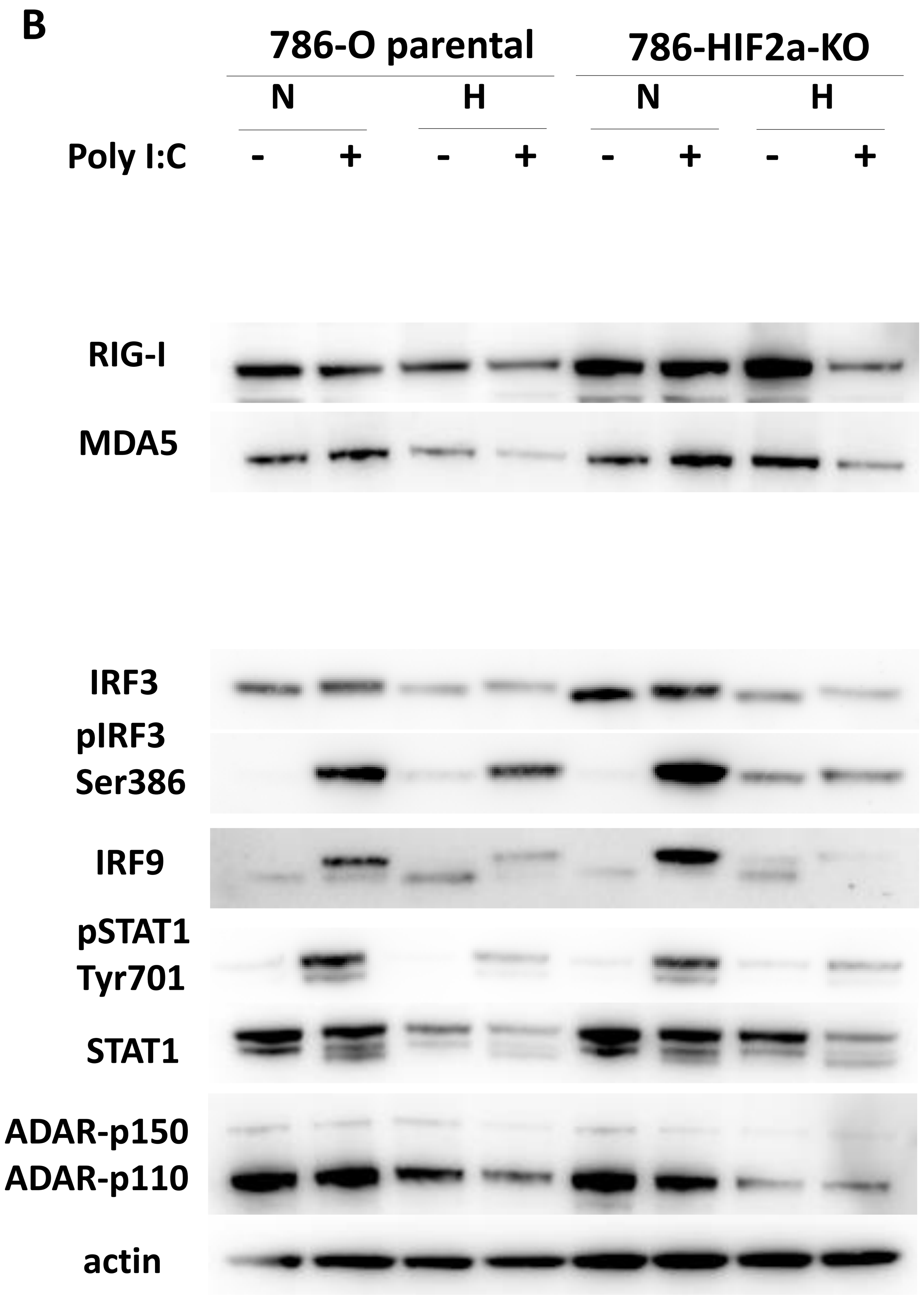


**Supplementary figure 3. HIF1α and HIF2α contribution to type I IFN downregulation in hypoxia**. Protein levels of type I IFN pathway members in RCC4-EV cells (HIF1α and HIF2α are upregulated even in normoxia due to a VHL mutation) and RCC4-VHL overexpressing cells (HIF1α and HIF2α are downregulated) cultured in normoxia or 0.1% hypoxia for 48h and transfected with poly I:C in the last 6h (n=3).
